# Alternative splicing and translation play important roles in parallel with transcriptional regulation during rice hypoxic germination

**DOI:** 10.1101/371583

**Authors:** Mo-Xian Chen, Fu-Yuan Zhu, Feng-Zhu Wang, Neng-Hui Ye, Bei Gao, Xi Chen, Shan-Shan Zhao, Tao Fan, Yun-Ying Cao, Tie-Yuan Liu, Ze-Zhuo Su, Li-Juan Xie, Qi-Juan Hu, Hui-Jie Wu, Shi Xiao, Jianhua Zhang, Ying-Gao Liu

## Abstract

Post-transcriptional mechanisms, including alternative splicing (AS) and alternative translation initiation (ATI), have been used to explain the protein diversity involved in plant developmental processes and stress responses. Rice germination under hypoxia conditions is a classical model system for the study of low oxygen stress. It is known that there is transcriptional regulation during rice hypoxic germination, but the potential roles of AS and ATI in this process are not well understood. In this study, a proteogenomic approach was used to integrate the data from RNA sequencing, qualitative and quantitative proteomics to discover new players or pathways in the response to hypoxia stress. The improved analytical pipeline of proteogenomics led to the identification of 10,253 intron-containing genes, 1,729 of which were not present in the current annotation. Approximately 1,741 differentially expressed AS (DAS) events from 811 genes were identified in hypoxia-treated seeds in comparison to controls. Over 95% of these were not present in the list of differentially expressed genes (DEG). In particular, regulatory pathways such as spliceosome, ribosome, ER protein processing and export, proteasome, phagosome, oxidative phosphorylation and mRNA surveillance showed substantial AS changes under hypoxia, suggesting that AS responses are largely independent of traditional transcriptional regulation. Massive AS changes were identified, including the preference usage of certain non-conventional splice sites and enrichment of splicing factors in the DAS datasets. In addition, using self-constructed protein libraries by 6-frame translation, thousands of novel proteins/peptides contributed by ATI were identified. In summary, these results provide deeper insights towards understanding the underlying mechanisms of AS and ATI during rice hypoxic germination.

## INTRODUCTION

Rice is a staple food that provides dietary nutrition for more than two billion people around the world (Yang and Zhang, 2006). In addition, rice is a model monocot plant used in modern research. Rice has been reported to have the ability to survive periods of submergence from seed germination to adult plants (Atwell et al., 2015). In particular, it has been documented as one of the few species that can germinate under anoxia by elongating its coleoptile to reach the water surface (Berta and Ismail, 2013). This adaptation to oxygen deprivation caused by flooding can be used as a model to study molecular mechanisms in response to hypoxic or anoxic conditions. Flooding is becoming one of the most severe abiotic stresses worldwide (Sasidharan et al., 2017). As the primary stresses due to flooding, hypoxia and anoxia have drawn much attention in the past decade. Under normal oxygen concentrations, oxygen gradients have been reported in dense plant organs, including seeds, fruits and tubers (Sasidharan et al., 2017). Thus, studying the molecular mechanisms during hypoxic or anoxic conditions may facilitate an understanding of the function of O_2_ molecules in both stress response and plant development. In recent years, a N-end rule protein degradation pathway has been proposed to be an important oxygen sensing mechanism in Arabidopsis (Gibbs et al., 2011; Licausi et al., 2011). Its downstream components, plant ethylene-responsive transcription factors, are affected by this pathway to activate or deactivate their target genes in response to hypoxia (Weits et al., 2014; Giuntoli et al., 2017). Increasing numbers of loci involved in flooding responses have been characterized, including those in lipid signalling (Xie et al., 2015), jasmonic acid and antioxidant pathways (Yuan et al., 2017), protein kinase (Chang et al., 2012) and transcription factor (Giuntoli et al., 2017). However, few studies are related to functional characterization of rice genes during flooding germination. In addition to the classical CIPK15-SnRK1A-MYBS1-mediated sugar-sensing pathway (Lu et al., 2007; Lee et al., 2009), a mitochondrion-localized protein (OsB12D1) has been reported to enhance flooding tolerance in rice germination and subsequent seedling growth (He et al., 2014). In addition, a rice trehalose-6-phosphate (T6P) phosphatase (OsTPP7) gene has been proposed to increase sink strength in response to flooding germination (Kretzschmar et al., 2015).

With the development of large profiling techniques, considerable efforts have been made to study the global transcripts changes, protein abundance and metabolic variation during rice hypoxic germination (Lasanthi-Kudahettige et al., 2007; Narsai et al., 2009; Narsai et al., 2011; Sadiq et al., 2011; Narsai et al., 2015; Hsu and Tung, 2017). Although comparative analysis indicates that part of the hypoxic responsive pathways are conserved among several species (Narsai et al., 2011), the mechanism of adult flooding tolerance may be largely different from that of seed flooding tolerance (Lu et al., 2007; Lee et al., 2009). Furthermore, recent RNA seq analysis using eight Arabidopsis ecotypes suggests that alternative splicing could be another pivotal factor involved in hypoxic responses (Van et al., 2016). Alternative splicing results from post-transcriptional control of eukaryotic intron-containing genes. Recent advancement reveals that more than 95% of genes have splicing isoforms in mammals (Eckardt, 2013). Two major types of splicing complex have been documented that can determine the splicing site sequences. One is the U2 complex, which can splice at a 5’-GT-AG-3’ exon-intron junction. The other is called the U12 complex and is able to utilize 5’-AT-AC-3’ as a splicing junction (Zdraviko J et al., 2005; Will and Luhrmann, 2011). Alternative splicing from multiexonic genes has been regarded as a potential way to increase plant genome coding ability (James et al., 2012; Ruhl et al., 2012; Chang et al., 2014; Feng et al., 2015). In addition to alternative splicing, another type of post-transcriptional regulation defined as alternative translation initiation (ATI) is involved in contributing to protein diversity (Sonenberg and Hinnebusch, 2009). Recent identification of translation initiation sites using advanced technology such as ribosome sequencing and MS-based proteomics reveals that a large number of these sites are not conventional AUG sequences (Sonenberg and Hinnebusch, 2009; Ingolia et al., 2011; Lee et al., 2012). In comparison to AS regulation (James et al., 2012; Ruhl et al., 2012; Chang et al., 2014; Feng et al., 2015; Wang et al., 2015; Zhan et al., 2015; Thatcher et al., 2016), the function of ATI has been seldom reported in plants (de Klerk and t Hoen, 2015). The above techniques have demonstrated that the eukaryotic genome has the ability to encode short peptides, including upstream open reading frames (uORFs) and other small ORFs, that are located in previously marked non-coding regions of the genome (Tavormina et al., 2015). Several peptides have been characterized to show crucial roles in regulating plant development and stress responses (Simon and Dresselhaus, 2015; Tameshige et al., 2016).

In summary, although stress-induced genome-wide AS changes have been extensively documented in various plant species (Yang et al., 2015; Thatcher et al., 2016; Van et al., 2016; Fesenko et al., 2017), the quantification of corresponding AS isoforms at the protein level have seldom been reported. In this study, a parallel RNA seq and proteomic approach defined as proteogenomic has been applied to achieve integrative analysis using both transcriptome and proteome data. Given our previous experience in ABA-regulated AS analysis (Zhu et al., 2017), we further improved our analytical pipeline for the determination of AS- and ATI-induced genome coding ability. The results from this study further expand our understanding of genome coding ability in rice seeds, suggesting an underlying regulatory network resulting from AS and ATI during rice hypoxic germination. Understanding this hidden network may facilitate the agricultural production of rice that is suitable for direct seeding systems and provide guidelines for improving hypoxic tolerance in other crop species.

## Materials and methods

### Plant material, growth conditions and hypoxic treatment

Seeds of *Oryza sativa* (Nipponbare) were surface-sterilized with 20% bleach and 0.05% Tween-20 before treatments. Seeds (~30-50 individuals) were placed on petri dishes with wet filter paper and then were transferred to air control or hypoxia conditions under complete darkness. The hypoxia treatment was carried out using the Whitley H35 Hypoxystation (Don Whitley Scientific Limited, UK) with 3% O_2_ level at 28°C. Seed samples were harvested at 6 h after treatments and used for further transcriptomic and proteomic analysis.

### Rice seed RNA extraction and RNA sequencing

Rice seed total RNA were ground in liquid nitrogen and extracted using a Plant RNeasy Mini Kit (Qiagen, Germany) according to the manufacturer’s instructions. RNA-sequencing (RNA-seq) experiments were conducted as previously described with minor modifications (Zhu et al., 2017). The resulting cDNA library constructed from rice seed RNA samples (Air_6 h and Hypoxia_6 h) were used for paired end (2 × 125 bp) sequencing on an Illumina HiSeq 4000 platform by Annoroad Gene Technology Co. Ltd. (Beijing, China). Three replicates for each sample were trimmed to obtain clean reads for subsequent analysis (Supplemental Table 1).

### Analysis of RNA sequencing and proteomic data

The rice (Nipponbare) reference genome annotation file (Oryza_sativa.IRGSP-1.0.32) was downloaded from the Ensembl website (http://www.ensembl.org/index.html). Clean reads mapping and subsequent bioinformatic analysis was as described previously (Zhu et al., 2017). The analytical pipeline is summarized in Supplemental Fig. 1. As mentioned previously (Zhu et al., 2017), significant changes of differentially expressed genes (DEG) (Supplemental Table 2) and differentially expressed alternative splicing events (DAS) (Supplemental Table 3) were determined as Log_2_FC > 2 and *q*-value (false discovery rate, FDR < 5%). Identification and quantification of AS events were conducted by using the software ASprofile (http://ccb.jhu.edu/software/ASprofile) (Florea et al., 2013). Splicing junctions reported in this study was generated by default settings of TopHat v2.1 aligner. The AS events with no expression values were filtered out before subsequent analysis (Zhu et al., 2017). Gene ontology analysis (GO, http://geneontology.org/) and Kyoto encyclopedia of genes and genomes (KEGG, http://www.kegg.jp/) enrichment classification were carried out using both DEG and DAS datasets. Heatmaps were generated using the BAR HeatMapperPlus tool (http://bar.utoronto.ca/ntools/cgi-bin/ntools_heatmapper_plus.cgi). The splicing sites conservation analysis was performed using WebLogo v3 (http://weblogo.threeplusone.com/) (Crooks et al., 2004).

### Total protein extraction, digestion and qualitative identification

Total protein of rice seeds was extracted and digested as described previously (Chen et al., 2014) with minor modifications. In general, approximately 5 g of rice seed tissues of each sample were ground in liquid nitrogen for subsequent proteomic analysis. The precipitated protein pellets were digested by trypsin and desalted using a Sep-Pak C_18_ column (Waters). The resulting peptides were then separated and characterized in a TripleTOF 5600^+^ (AB SCIEX) splitless Ultra 1D Plus (Eksigent) system (Andrews et al., 2011).

### Peptide dimethyl labelling and quantitative proteomics

The quantitative proteomics was conducted as described previously with minor modifications (Zhou et al., 2015). Digested peptides were dissolved with 0.1 M sodium acetate (pH≈6, best below 6) (*i.e.*, 500 μg peptides per 0.25 mL sodium acetate). Either 4% formaldehyde or formaldehyde-d2 (40 μL per 500 μg peptides) were added and mixed. Then, 40 μL / 500 μg peptides of 0.6 M NaBH_3_(CN) were added. The solution mixture was shaken for 0.5 h. Furthermore, 160 μL / 500 μg peptides of 1% NH_4_OH, was added and mixed for 5 min. Then, 5% formic acid (160 μL per 500 μg peptides) was added and mixed. The solution was placed in 4°C for at least 1 h. The light and heavy dimethyl labelling peptides were combined in a 1:1 ratio and desalted using a Sep-Pak C_18_ column (Waters).

Mixed peptides were subsequently fractionated by using a C_18_-ST column (2.0 mm × 150 mm, 5 μm particle size) (TechMate) on the Agilent 1260 system (Agilent Technologies). An elution gradient of 60 min was used for peptide separation with 20 mM ammonium formate in H_2_O (adjusted to pH 10 by 25% NH_3_.H_2_O) as solvent A and 20 mM ammonium formate in 80% ACN (adjust pH to 10 by 25% NH_3_.H_2_O) as solvent B. The gradient elution profile was composed of 5%-25% B for 20 min, 25-45% B for 15 min, 45-90% B for 1 min, then maintained at 90% B for 4 min, followed by 10-95% A for 1 min, and ending with 95% A for 14 min. The flow rate was 0.2 mL/min. UV absorbance was monitored at 216 nm. A total of 60 0.2 mL fractions were collected, then concatenated and mixed to obtain 20 fractions. Fractions were dried *via* speed-vacuum and desalted by the StageTip C_18_ method. RPLC-ESI-MS/MS was used to detect the sample. LC-MS/MS detection was carried out on a hybrid quadrupole-TOF LC/MS/MS mass spectrometer (TripleTOF 5600^+^, AB Sciex) equipped with a nanospray source. Peptides were first loaded onto a C_18_ trap column (5 μm, 5 x 0.3 mm, Agilent Technologies) and then eluted into a C_18_ analytical column (75 μm × 150 mm, 3 μm particle size, 100 Å pore size, Eksigent). Mobile phase A (3% DMSO, 97% H2O, 0.1% formic acid) and mobile phase B (3% DMSO, 97% ACN, 0.1% formic acid) were used to establish a 100 min gradient, which consisted of 0 min of 5% B, 65 min of 5-23% B, 20 min of 23-52% B, 1 min of 52–80% B, and the gradient was maintained in 80% B for 4 min, followed by 0.1 min of 80–85% B, and a final step in 5% B for 10 min. A constant flow rate was set at 300 nL/min. MS scans were conducted from 350 to 1500 amu, with a 250 ms time span. For MS/MS analysis, each scan cycle consisted of one full-scan mass spectrum (with m/z ranging from 350 to 1500 and charge states from 2 to 5) followed by 40 MS/MS events. The threshold count was set to 120 to activate MS/MS accumulation, and former target ion exclusion was set for 18 s.

### Library construction and mass spectrometry database searching

An AS junction library (576,570 entries) was constructed as described previously (Sheynkman et al., 2013; Castellana et al., 2014; Walley and Briggs, 2015) with minor modifications. In brief, six-frame translations, including 3 frames on the forward strand and 3 frames on the reverse complement strand, were used construct the AS junction library. Additionally, a frame library was constructed using all transcripts annotated in the reference annotation file by 6 frames. The redundant sequences were then removed from translated sequences at the first step. Peptide sequences longer than 6 amino acids were attached to the UniProt rice japonica database for subsequent database search. Raw spectrum data generated from both qualitative and quantitative proteomics were searched with the ProteinPilot software (v5.0, AB SCIEX) using preset parameters. All data were filtered at 1% FDR with at least 1 peptide at 95% confidence level calculated automatically by the ProteinPilot software (Zhu et al., 2017). For quantitative proteomics, data were searched against UniProt and self-constructed databases using the following parameters: sample type, dimethyl (0,+4) quantitation; cys alkylation, iodoacetamide, digestion, trypsin. The search effort was set to rapid ID. For DEP analysis, proteins with a fold change of >1.2 or <0.8 (*P* value <0.05) are considered as DEP in this study.

### Quantitative real-time PCR validation of AS transcripts

Total RNA (~5 μg) was reverse-transcribed into cDNA by using the Superscript First-Strand Synthesis System (Invitrogen, USA) following the manufacturer’s instructions. Quantitative real-time PCR (qRT-PCR) was conducted as described previously (Zhu et al., 2013). Resulting products of qRT-PCR were subjected to DNA sequence analysis. Isoform-specific primers used for AS isoforms identification are listed in Supplemental Table 5.

### Data submission

The rice transcriptome data have been uploaded to Sequence Read Archive (https://www.ncbi.nlm.nih.gov/sra) under Bioproject PRJNA451248. The raw data of qualitative and quantitative proteomics have been submitted to the PRIDE PRoteomics IDEntifications (PRIDE) database with accession number PXDxxxxxx and PXDxxx, respectively.

## Results

### Improvement of analytical pipeline and experimental conditions

The analytical pipeline used in this study is presented in Supplemental Fig. 1. Improvements have been made since the last bioinformatic flowchart (Zhu et al., 2017). The identification and quantification procedures of AS events were simplified for subsequent GO and KEGG analysis. In addition, refinement of redundancy and error check steps further improved the accuracy of identification. In this study, AS events such as AFE (alternative first exon) and ALE (alternative last exon) purely caused by alternative transcription start and poly adenylation has been removed to further differentiate AS modification from other transcriptional or post-transcriptional mechanisms. To distinguish 5’ donor sites and 3’ acceptor sites, we further divided AE (alternative exon) events into AE5’ and AE3’ for further bioinformatic analysis. Furthermore, incorporation of quantitative proteomics yielded more information on steady protein levels in comparison to qualitative proteomic profiling, which can only identify the presence of translated peptides (Zhu et al., 2017). For testing samples, we chose dry seeds of japonica rice (Nipponbare) treated with hypoxia (3% O_2_) for 6 h in comparison to air controls under complete darkness. This treatment will help us to understand the short-term responses at both transcripts and protein levels during hypoxia when seeds start to germinate. Plenty of samples were harvested for the following three profiling experiments: short-read RNA sequencing (RNA seq), qualitative and quantitative proteomics. Prior to these experiments, we have compared 49 up-regulated anaerobic marker genes highlighted in previous publications to our dataset (Lasanthi-Kudahettige et al., 2007; Narsai et al., 2009). Among 21 genes detected in this study, 19 of these genes showed consistency on their differential regulation, but at a lower magnitude (Supplemental Fig. 2A). We used qRT-PCR to further validate those expressions. In total, 18 of 19 genes showed similar expression pattern as the result of our RNA seq data (Supplemental Fig. 2B), indicating the efficacy of hypoxic treatment using 3% O_2_ in our system.

### Completely different set of genes undergo alternative splicing (AS) in response to hypoxia during rice seed germination

Approximately 1.32 billion raw reads in total averaging 200 million reads per sample were obtained from RNA sequencing (Supplemental Table 1). Among these, 1.25 billion clean reads were subjected to the mapping process. On average, approximately 95% were uniquely mapped to the genome and used for subsequent bioinformatic analysis (Supplemental Table 1). For AS identification, each sample identified over 75,000 AS events. In total, 10,253/26,848 (38.2%) annotated intron-containing genes in rice seed were observed to exist as AS events in rice seeds. Approximately 6.4% (1,729/26,848) more intron-containing genes were observed in comparison to the original annotation file. Slightly differ from previous AS analysis in ABA-treated Arabidopsis seedlings (Zhu et al., 2017), alternative first exons (AFE), alternative last exons (ALE) and intron retention (IR) remained as the most abundant three AS events through all the samples (Fig. 1A). Among these three AS event types, AFE and ALE caused variable 5’- and 3’-untranslated ends, which may affect the efficiency of translation or stability of corresponding transcripts (Andreassi and Riccio, 2009; Sonmez et al., 2011; Jenal et al., 2012). For example, hidden small open reading frames (sORF) from the 5’-end of transcripts encoding short peptides have the ability to regulate translational efficiency of target transcripts (Laing et al., 2015), whereas polyadenylation at the 3’-end of transcripts is well known to affect the localization and stability of the transcripts (De et al., 2017). When the dataset of differentially expressed genes (DEG) (Supplemental Table 2) was compared to the dataset of differentially expressed AS genes (DAS) (Supplemental Table 3), over 95% were not the same (Fig. 1B). Only 23 genes were differentially regulated at both transcription and post-transcriptional levels (Fig. 1B). This suggests that alternative splicing may play an important and distinctive role during rice hypoxic germination. Subsequent gene ontology enrichment analysis also confirmed the result from the Venn diagram (Fig. 1B, Supplemental Fig. 3). In several cases, DEG and DAS genes did not coexist in the same secondary GO category (Supplemental Fig. 3). Fourteen isoforms of 7 genes in the DAS dataset were assembled and validated by quantitative real-time PCR (qRT-PCR). In total, 6 of these genes were consistent to the data from RNA seq analysis, suggesting the reliability of AS identification and quantification from the analytical pipeline (Supplemental Fig. 4). Except for categories related to linoleic acid metabolism, the majority of DEG and DAS genes were not enriched in the same KEGG category (Supplemental Fig. 3), suggesting that DAS category is a different group of genes in response to hypoxic germination. The majority of pathways enriched in DEG dataset were closely related to cellular metabolisms (*e.g.* pentose phosphate pathway, glycolysis/gluconeogenesis, fructose and mannose metabolism *etc.*) and cell growth (meiosis, DNA replication and cell cycle *etc.*). Whereas some regulatory pathways were specifically over-represented in DAS dataset, such as spliceosome, ribosome, ER protein processing, protein export, proteasome, phagosome, oxidative phosphorylation and mRNA surveillance pathway, implying that these pathways may play essential role in AS-mediated responses under rice hypoxic germination. Gene members in several pathways have been selected for RT-PCR and qRT-PCR validation (Fig. 1D and Fig. 2). Some splicing isoforms of corresponding gene showed differential expression under hypoxic treatment, indicating their potential role in response to rice hypoxic germination.

**Figure 1.**
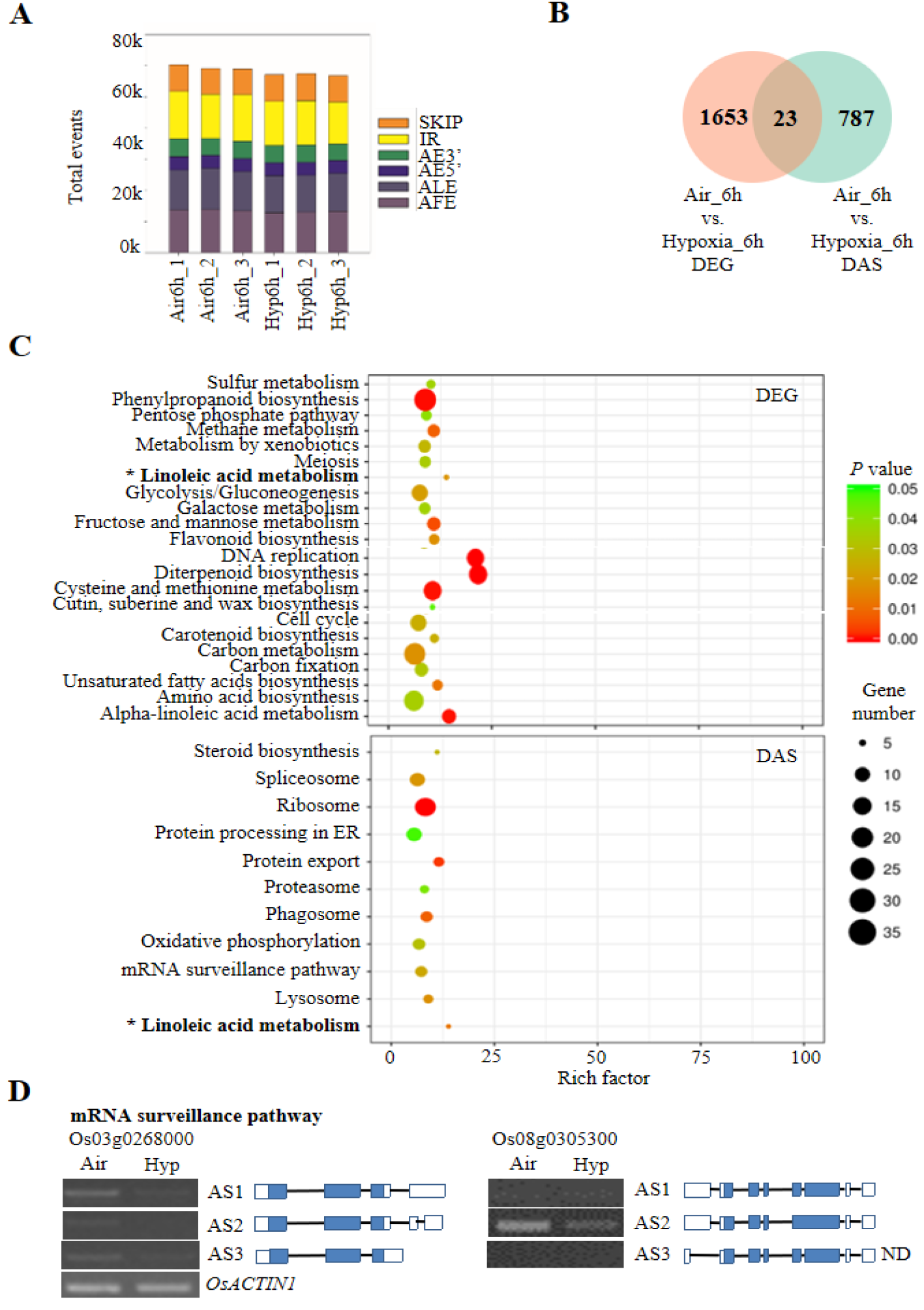
Identification and comparison between the datasets of differentially expressed genes (DEG) and differentially expressed alternative splicing (DAS) events during rice hypoxic germination. (A) Statistics of the identified alternative splicing (AS) events and types. ALE, alternative last exon; AFE, alternative first exon; SKIP, exon skipping; IR, intron retention; AE5’, alternative donor; AE3’, alternative acceptor. (B) The Venn diagram represents unique and shared genes between DEG and DAS datasets. (C) Gene ontology enrichment analysis between DEG and DAS datasets. (D) RT-PCR validation of the DAS events in mRNA surveillance pathway. Air: air control; Hyp: Hypoxia; ND: not detected. Gene models of each isoform are indicated (blue: coding region; white: non-coding UTRs; not to scale).

**Figure 2.**
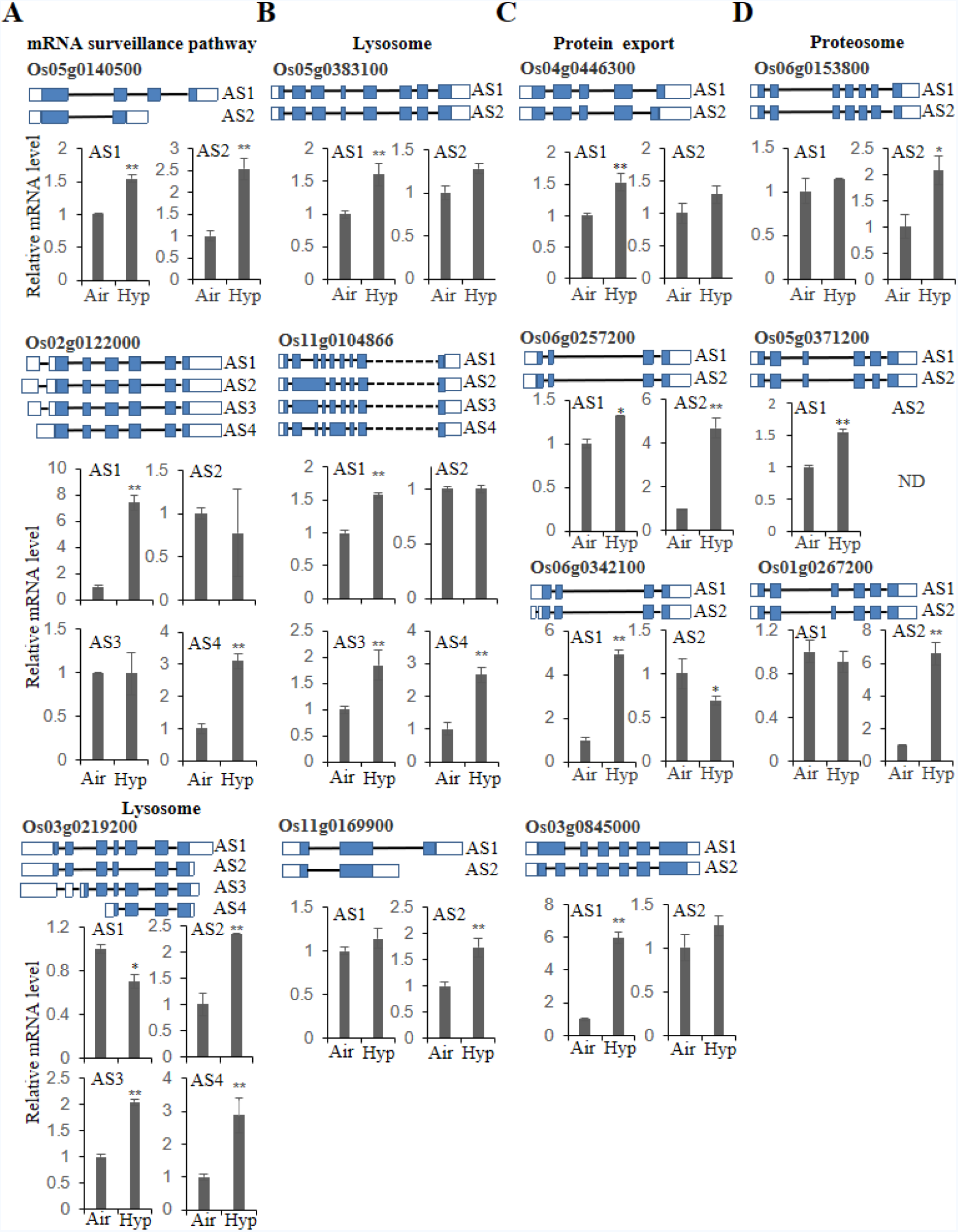
qRT-PCR validations of DAS events. Validation of DAS events detected in KEGG enrichment analysis. DAS events involved in categories of (A) mRNA surveillance pathway and lysosome, (B) lysosome, (C) protein export, (D) proteasome was verified by qRT-PCR analysis from three biological replicates. *OsACTIN1* was used as an internal reference gene. Mean values±SD are presented (n=3). ‘**’ and ‘*’ represent mean values of hypoxia-treated (Hyp) group is significantly higher or lower than that in air control (Air), *P*<0.01 and *P*<0.05, respectively. Gene models of each isoform are indicated (blue: coding region; white: non-coding UTRs; not to scale).

### Qualitative proteomic identification reveals that hypoxia-regulated AS events are more likely to be translated

To further characterize the translational products of identified AS events, we carried out a qualitative proteomic profiling using tandem mass spectrometry (MS/MS) for both control and hypoxia-treated samples (Alfaro et al., 2014; Tavares et al., 2015; Zhu et al., 2017). Proteomic analysis this time generated 547,545 and 485,392 high-quality spectra for control and hypoxia-treated samples, respectively. Approximately 5,549 and 5,385 proteins were identified using the UniProt database (Fig. 3A). Among these, 18.6% and 16.1% of identified proteins were uniquely present in control or hypoxia-treated samples, respectively, serving as good candidates for further functional characterization. Subsequent AS junction library search identified 4,431 / 4,313 peptides from AS events (41,887) and 510 / 490 peptides from DAS events (1,742) for control / hypoxia-treated samples, respectively (Fig. 3A, B). Among these, approximately 70% of peptides were shared by both samples. Intriguingly, much fewer AFE events could be detected at peptide level in comparison to ALE events (Fig. 2B). Furthermore, 13.5% of the total AS events (5,652/41,887) were translated into peptides, suggesting that the majority of AS transcripts may be degraded by RNA surveillance mechanisms such as nonsense-mediated mRNA decay (NMD) (Nicholson et al., 2010; Drechsel et al., 2013). In contrast, an elevated percentage (38.3%) of DAS events could be translated into peptides in all AS types (Fig. 3B), indicating their potential role in response to hypoxic stress during rice germination. Similar observations have been reported in ABA-treated Arabidopsis seedlings, which indicates that thousands of AS proteins are translated under hypoxic conditions during rice germination, and most of these were not present in the DEG dataset analysed by a conventional RNA seq pipeline.

**Figure 3.**
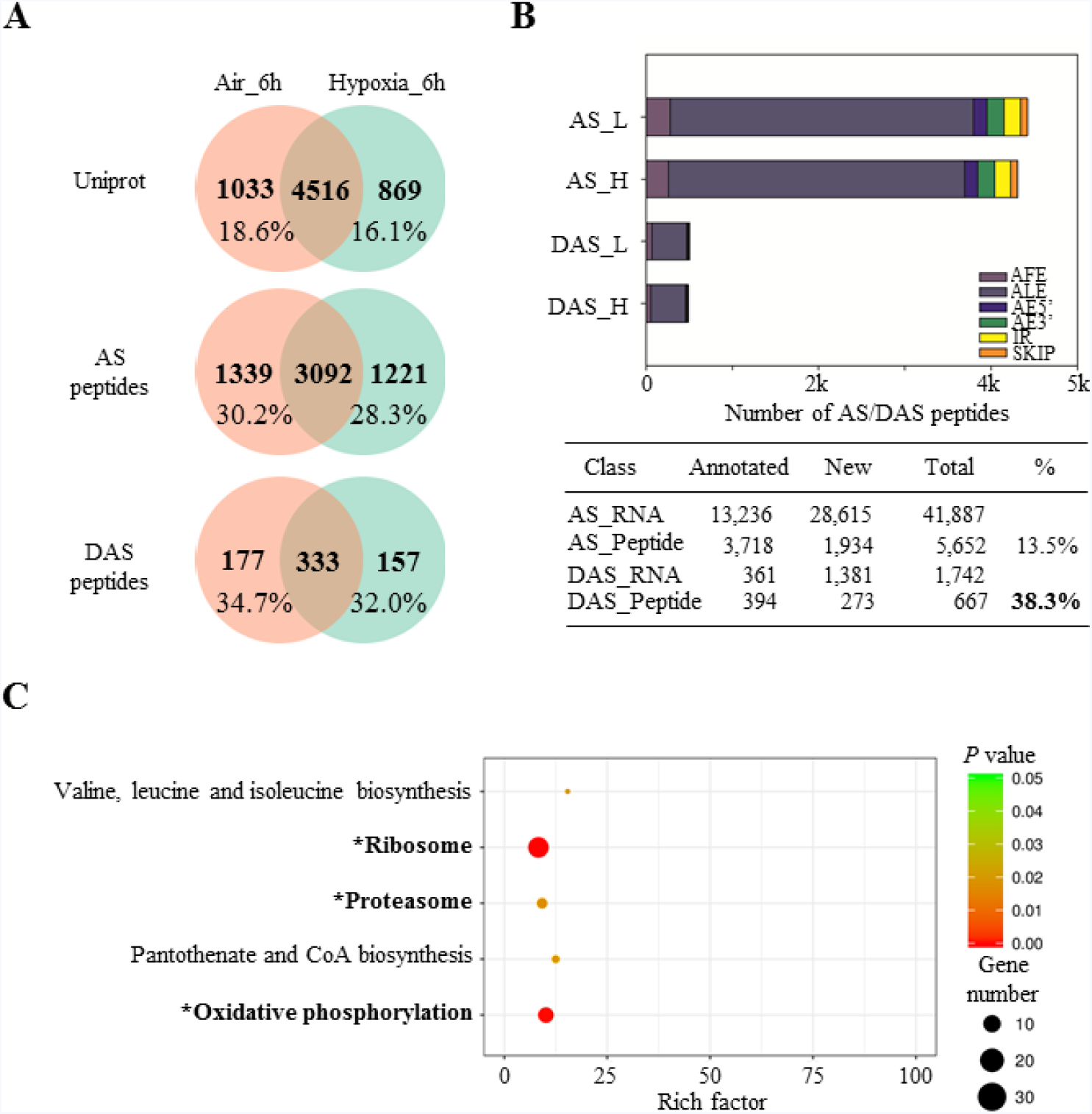
Qualitative proteomic identification of AS peptides. (A) Venn diagram representation of qualitative proteomic identification using UniProt, AS and DAS databases. (B) AS peptides identification and classification (upper panel). ALE, alternative last exon; AFE, alternative first exon; SKIP, exon skipping; IR, intron retention; AE5’, alternative donor; AE3’, alternative acceptor. Summary of identified AS/DAS events and peptides (lower panel). (C) KEGG pathway enrichment analysis of DAS peptides in qualitative proteomics. ‘*’ marked pathway is repeatedly found in both transcriptome and qualitative proteomic datasets.

In addition, approximately 68.3% of AS events identified in this study were not annotated in the genome and thus were marked as new features for rice genome annotation (Fig. 3B). Additionally, 40.9% of the DAS peptides were not present in the current version of the annotation, which suggests the translation of new protein isoforms during rice germination in response to hypoxia. DAS peptides were subjected to KEGG enrichment analysis (Fig. 3C). For example, some KEGG terms including ribosome, proteasome and oxidative phosphorylation, were repeatedly enriched in both RNA seq and qualitative proteomic datasets, giving protein evidence of these splicing isoforms in response to hypoxia treatment.

### Quantitative proteomics indicates that the expression of protein and transcripts are correlated at the AS level

To find relationship between the protein abundance and corresponding transcripts at the AS level, quantitative proteomics were conducted using the dimethyl labelling method. In total, 10,946 proteins were identified from this approach and 4566 of them were quantified (Supplemental Table 4). Among these, 278 differentially expressed proteins (DEP) and 29 differentially regulated AS peptides (DASP) were identified (Fig. 4A-B). Thirteen DASP were found to be differentially expressed in quantitative proteomics and referred as DASDP. Amongst these, none of them were shared with DEP dataset (Fig. 4C). Similar to previous parallel analysis (Bai et al., 2015; Marmiroli et al., 2015), much less overlap was observed between DEP and DEG, DAS and DASP as well as DEP and DASDP (Fig. 4A-C). Only 11 genes were identified as both DEG and DEP with low correlation (R^2^=0.18) of their expression levels (Fig. 4D, E), suggesting the existence of post-transcriptional regulation for most of the transcripts. Although 2 genes were detected in both DAS and DASP datasets, the expression of their transcripts and proteins were at the same trend (Fig. 4F), indicating that quantification at AS isoform level may provide more accurate data representation for both transcripts and proteins than conventional quantification method used in RNA seq and proteomics. However, more data is required to confirm this hypothesis. In addition to the effect of post-transcription, the low overlap of DEP/DASDP with DEG/DAS datasets may be explained by the relatively low throughput and coverage of the MS-based proteomic method.

**Figure 4.**
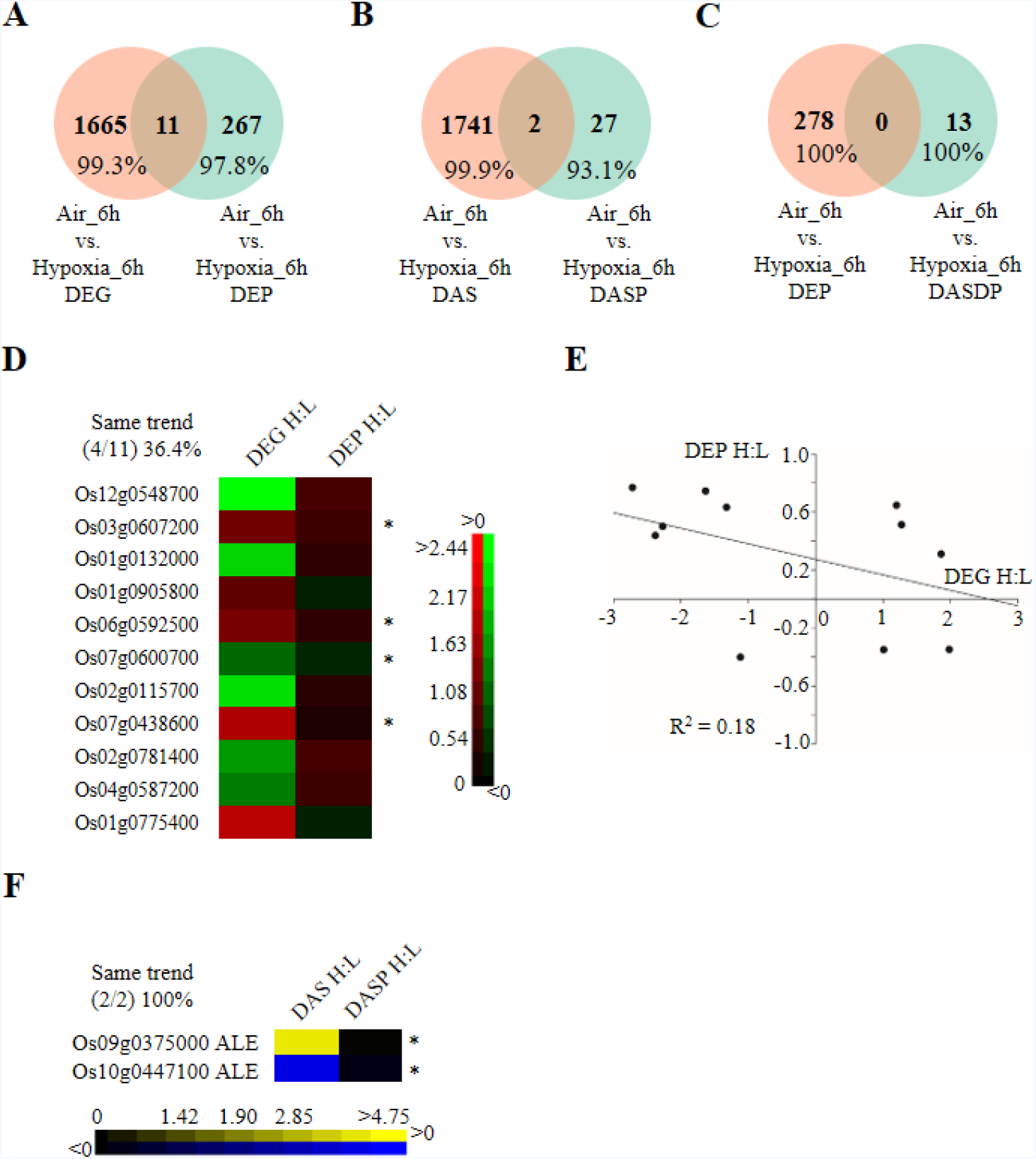
Comparison between proteomic and transcriptomic datasets. Venn diagram representation of (A) differentially expressed genes (DEG) vs. differentially expressed proteins (DEP), (B) differentially expressed AS events (DAS) vs. differentially expressed AS peptides (DASDP), (C) DEP vs. DASDP. Heatmap representation (D) and correlation analysis (E) of overlapping genes between DEG and DEP. (F) Heatmap representation of overlapping genes between DAS and DASDP. ALE, alternative last exon; ‘*’ represents the regulation of transcripts and proteins at the same trend in corresponding datasets; H: L, hypoxia vs. air control.

### Construction of a customized protein library leads to novel proteins identification and quantification during rice hypoxic germination

Similar to previous findings (Zhu et al., 2017), the spectra usage for protein identification was approximately 40-50% in this study (Fig. 5A) using both UniProt and AS junction libraries as input files. An increasing number of publications suggest that single transcripts are able to be translated into multiple proteins by using alternative translation initiation (ATI) sites (Brar and Weissman, 2015). This indicates that a large number of novel proteins or short peptides are yet to be identified, and this is caused by incomplete genome annotation (Kim et al., 2014). Thus, a 6-frame translation library was constructed using the combination of assembled cufflink files during RNA seq analysis and reference annotation files based on previously published methods (Castellana et al., 2008; Zhu et al., 2017). The outcomes from the database searching identified thousands of novel proteins and peptides, with 74.6% of proteins longer than 80 amino acids (a. a.), 24.0% of proteins/peptides from 11-80 amino acids and 1.4% of peptides from 6-11 amino acids (Fig. 5B, C). Among these, 2294 / 1432 novel proteins (> 80 a.a.) and 310 / 774 novel proteins or peptides (6-80 a. a.) were identified in control / hypoxia-treated samples, respectively (Fig. 5D). This observation provides further evidence of increment coding ability for proteins and short peptides by using ATI sites. Additionally, an increasing number of short peptides (774) were detected in hypoxia-treated samples in comparison to air controls (310), suggesting that short peptides may play an important role in response to hypoxia during rice germination. Intriguingly, 137 novel proteins were quantified at a second frame of known transcripts. Few of these overlapped with DEG and DEP datasets, indicating that most of these proteins can only be detected by proteomic analysis using the customized library. This set of genes served as a source of novel candidates for further investigation of hypoxic responses during rice germination.

**Figure 5.**
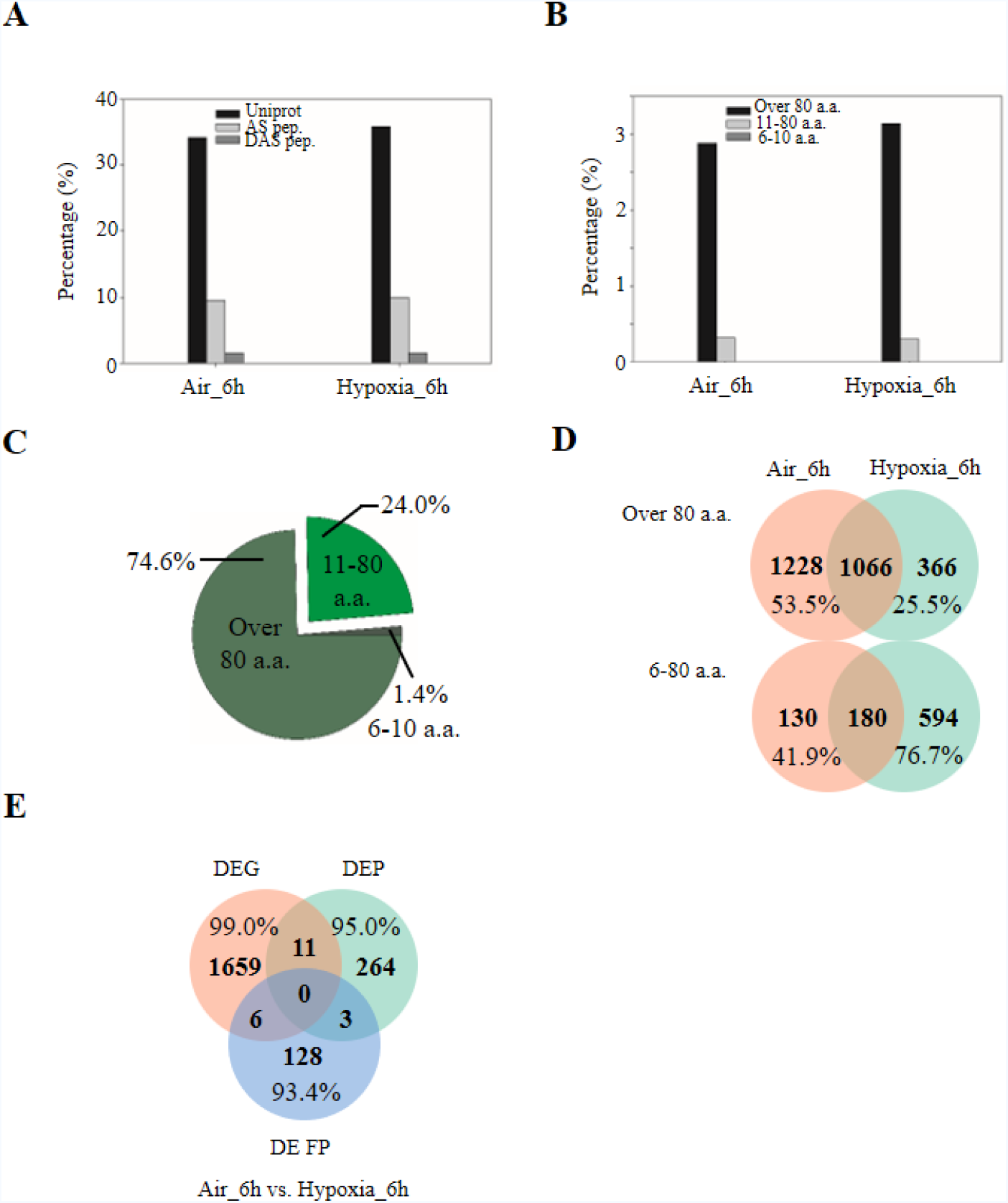
Novel protein/peptide identification. (A) Summary of the spectrum usage of the data from qualitative proteomics using UniProt, AS and DAS databases. (B) Spectrum usage of the data from qualitative proteomic using the 6-frame translated protein database. (C) Pie diagram represents percentage distribution of identified novel proteins/peptides. (D) Venn diagram representation of identified novel proteins/peptides in control and hypoxia-treated samples. (E) Venn diagram represents the shared and unique genes among differentially expressed genes (DEG), differentially expressed proteins (DEP) and differentially expressed frame proteins (DE FP).

### The conventional 5’-splicing sites are less conserved in rice seeds at normal condition and under hypoxia treatment

To further investigate the splicing characteristics between total AS and hypoxia-affected DAS datasets, statistical analysis of splicing sites (ss) conservation was performed. Conventionally, U2-type splicing sites (5’-GT-AG-3’) are conserved and account for 90% of total splicing sites among plant species (Will and Luhrmann, 2011). In this study, the 3’-splicing site (AG) was relatively conserved and accounted for over 80% in both control and hypoxia-treated samples (Fig. 6A). An extra ‘C’ was identified as a conserved sequence in both AS and DAS datasets (Fig. 6B). Thus, 3’-splicing sites were identified as ‘CAG’ in rice seeds, and the hypoxia treatment did not change this signature (Fig. 6B). However, there was a decrease in the ‘AG’ proportion in hypoxia-treated samples, which was associated with the increase in the proportions of several other ss sequences especially ‘AC’. In contrast, the conventional 5’-splicing site (GT) accounted for only 50% of total AS and was increased to approximately 60% in the hypoxia-affected DAS dataset (Fig. 6A). Meanwhile, non-conventional 5’-splicing sites such as ‘AA’ and‘CT’ was was largely reduced in the DAS dataset by comparing with AS dataset, suggesting its role in response to hypoxia stress (Fig. 6A). In addition, similar results were obtained by conservation analysis; ‘GGT’ signature was obtained in both AS and DAS datasets (Fig. 6B). Further investigation of ss among AS types demonstrated that AFE was responsible for the ‘GT’ reduction in both AS and DAS datasets (Supplemental Fig. 5). Although 3’-ss were more conserved, certain types of non-conventional splicing sites were induced among the specific AS types in the DAS dataset in comparison to the AS dataset such as 3’-TG and 3’-TT in AE5’, 3’-AC and 3’-GG in AE3’, 3’-GC and 3’-TG in IR and 3’-GC in SKIP (Supplemental Fig. 5). This result indicates that the AS regulation under hypoxia stress may be caused by alternative recognition of sequence of splicing sites.

**Figure 6.**
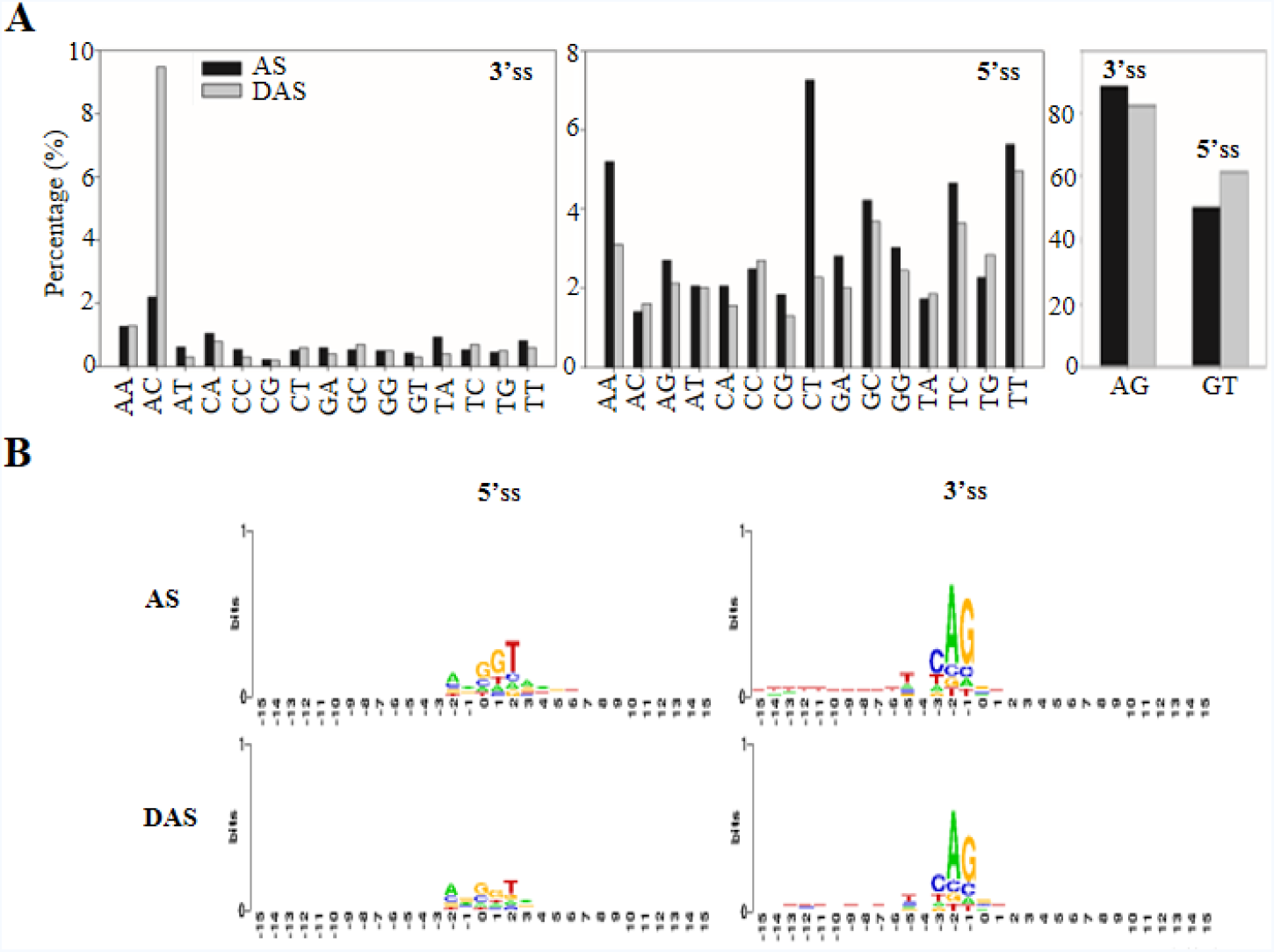
Splicing sites recognition under hypoxia stress. (A) Statistical analysis of the splicing sites (ss) between the total AS events and the hypoxia-affected DAS events. (B) Conservation analysis using sequence located at exon-intron junctions.

### Splicing factors are enriched in differentially expressed AS events

To further understand the underlying mechanism of DAS regulation under hypoxia stress, splicing factors in rice were summarized and subjected to further analysis. Three genes were found in the DEG dataset (Fig. 7A). In contrast, a total of 105 AS events from 21 splicing factor-related proteins were observed in the DAS dataset (Fig. 7A, B), and none of them were found in the DEG dataset. Amon these, 60 AS events were up-regulated, whereas 45 AS events were down-regulated (Fig. 7B). In detail, 43.8% of AS events were AFE and ALE accounting for 28.6% (Fig. 7C). The remaining three AS types accounted for 27.7% of the total AS events (Fig. 7C). According to the classification in the splicing-related gene database (SRGD, http://www.plantgdb.org/SRGD/index.php), the 21 genes observed in the DAS dataset were classified into 11 subgroups (Fig. 7D) from core splicing components to auxiliary factors. And those SFs enriched in KEGG term of spliceosome (Fig. 1C) were chosen for qRT-PCR validation (Fig. 7E). Some isoforms of selected SFs were differentially expressed under hypoxia treatment, suggesting that the change of AS in splicing components may be crucial in response to hypoxia stress during rice germination.

**Figure 7.**
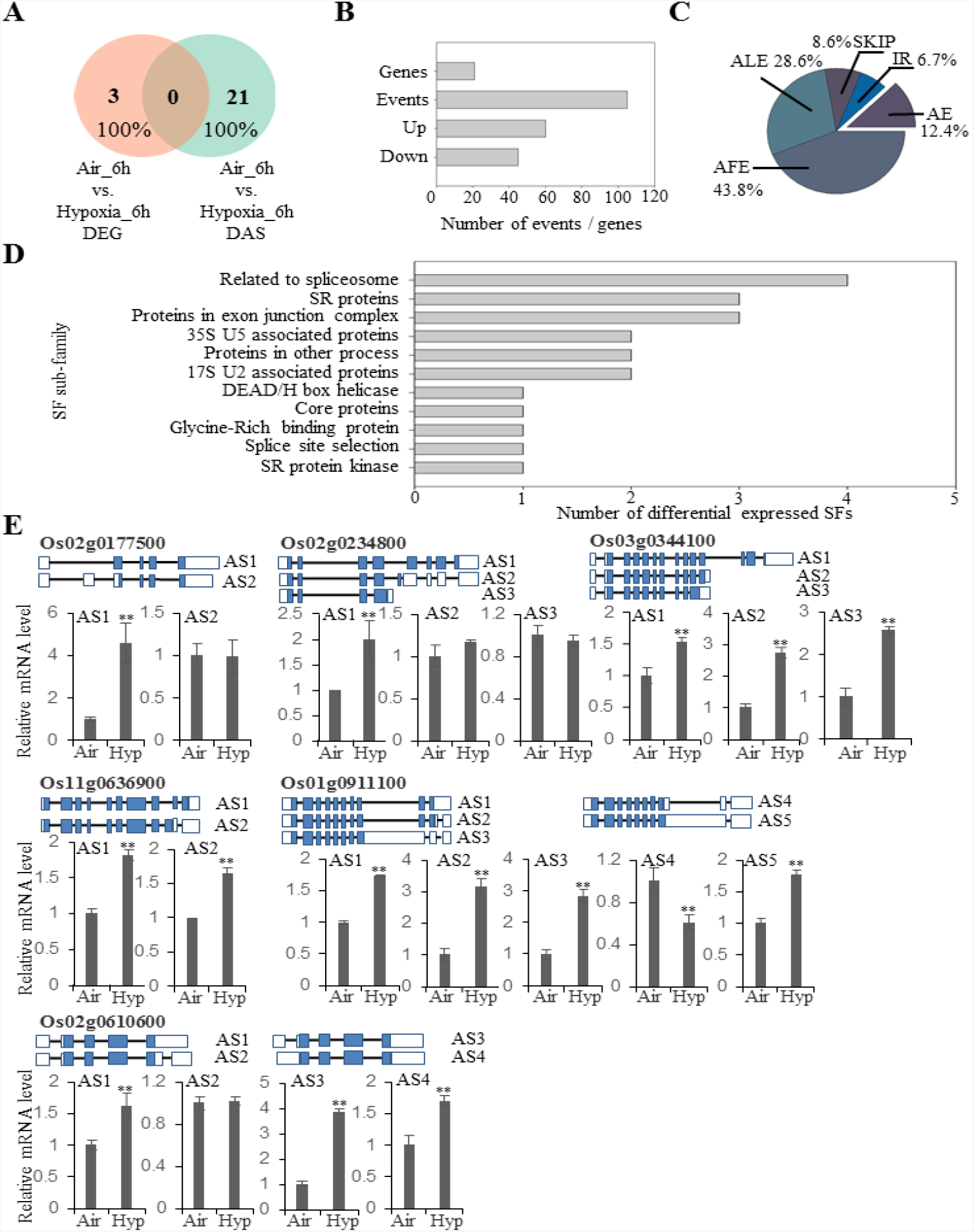
Splicing factors involved in hypoxia responses during rice germination. (A) The Venn diagram represents identified splicing factors between DEG and DAS datasets. (B) Statistics of hypoxia-affected DAS genes and events of splicing factors. (C) Pie chart distribution of the DAS events belonging to splicing factors. (D) Subgroup classification of splicing factors identified in DAS events. (E) qRT-PCR validation of DAS events detected in spliceosome. DAS events in the category of spliceosome were verified by qRT-PCR analysis from three biological replicates. *OsACTIN1* was used as an internal reference gene. Mean values±SD are presented (n=3). ‘**’ and ‘*’ represent the mean values of hypoxia-treated (Hyp) group is significantly higher or lower than that in air control (Air), *P*<0.01 and *P*<0.05, respectively. Gene models of each isoform are indicated (blue: coding region; white: non-coding UTRs; not to scale).

## Discussion

### The discovery of a hidden network of AS in response to hypoxic stress during rice germination provides additional targets in the study of hypoxia

Alternative splicing produces multiple RNA isoforms for each locus. Each isoform may encode one protein isoform as well, which greatly expands the genome coding ability. Additionally, the discovery of two new AS types, AFE and ALE, has revealed the great potential to generate AS isoforms (Yan and Marr, 2005; de Klerk and t Hoen, 2015; Zhu et al., 2017). In this study, approximately 97.2% (787/810) of the DAS genes had no differences at the gene expression level, suggesting AS control of transcripts is completely separated from the conventional DEG group (Fig. 1B). Moreover, increasing evidence reveals that protein isoforms generated by AS transcripts have the ability to alter protein subcellular localization, protein-protein interaction networks and protein stability due to the presence or absence of certain motifs (Buljan et al., 2012; Ellis et al., 2012). Thus, the 667 AS peptides identified in both control and hypoxia-treated samples provided protein evidence of AS transcripts and may serve as good candidates for further functional characterization (Fig. 3A, B). These genes were distributed in a variety of biological pathways, including amino acid biosynthesis, ribosome and proteasome pathway, pantothenate and CoA biosynthesis and oxidative phosphorylation, and were not selected for further investigation by the first round of screening using DEG as criteria amongst large scale transcriptome analysis, suggesting that AS responses are embedded in various biochemical processes under hypoxia stress.

Transcriptomic studies have shown that low oxygen induces a myriad of gene responsiveness in terms of transcript abundance (Lasanthi-Kudahettige et al., 2007; Narsai et al., 2009; Narsai et al., 2011; Sadiq et al., 2011; Narsai et al., 2015; Hsu and Tung, 2017). Accordingly, transcriptional regulation in oxygen sensing pathways has been extensively studied in plants. Key regulators, such as ERFVII transcription factors, have been substantially characterized (Fukao et al., 2009; Hattori et al., 2009; Hinz et al., 2010; Licausi et al., 2010). However, few studies have been carried out to unravel the AS regulation under hypoxia. In the current study, AS analysis indicates that the conventional splicing sites are not conserved at the 5’ position in rice seeds (Fig. 6A, B and Supplemental Fig. 5). Major regulators defined as splicing factors within assembled spliceosome have been characterized to participate in AS site determination (Golovkin and Reddy, 1996; Kalyna et al., 2006; Krummel et al., 2009; Will and Luhrmann, 2011; Kondo et al., 2015; Yoshida et al., 2015). Although several splicing factors have been reported to be involved in stress responses (Ruhl et al., 2012; Feng et al., 2015), none of them are related to hypoxia responses. In our results, a 7-fold increase in the number of splicing factors (21 in DAS to 3 in DEG) were found in comparison to the DEG dataset (Fig. 7), suggesting the importance of those proteins in splicing site recognition. Over 100 AS events in these 21 splicing factors were affected during hypoxia treatment, which may greatly alter the protein isoforms of these proteins in comparison to the control group. Subsequently, hypoxia may change the composition and conformation of spliceosomes by recruiting different protein isoforms of splicing factors, which may in turn lead to a different choice of splicing site sequence recognition. This may explain the increment of the proportion of certain non-conventional splicing sites during rice hypoxic germination (Supplemental Fig. 5). Furthermore, the integration of qualitative proteomic data implies that hypoxia-responsive AS events are more likely to be translated in comparison to non-responsive ones (Fig. 3B, lower panel), providing protein evidence for the potential role of these AS isoforms in response to hypoxia stress. Therefore, our results suggest that alternative splicing is an independent pathway other than transcriptional repression in response to hypoxia during rice germination. The majority of members in this pathway remain to be elucidated.

### Alternative cellular pathways are activated by AS under hypoxia treatment

Several pathways were found to be over-represented under AS-mediated responses during rice hypoxic germination. mRNA surveillance, such as NMD, has long been demonstrated to play an important role in controlling mRNA stability and abundance before translation (Nicholson et al., 2010; Drechsel et al., 2013). It has been reported that NMD is closely related to exon junction complex (EJC) of splicing machinery in both animals and plants (Shaul, 2015). In Arabidopsis, hypoxia-responsive ERFs, HRE1 and HRE2, have been proposed to be likely regulated by post-transcriptional mechanisms for their mRNA stability (Licausi et al., 2010). From our dataset, isoforms of several components belong to EJC complex (*e.g.* Os08g0305300, *OsSMG7* and Os05g0140500, *OsY14a*) were observed to be differentially regulated (Nyikó et al., 2013), indicating their potential function in surveillance of newly spliced RNA isoforms under hypoxia. Evidence shows that the status of spliceosome will be affected under hypoxia in animal tissues (Schmidtkastner et al., 2008). Splicing factors like serine-arginine (SR) proteins is activated under hypoxic condition by phosphorylation (Jakubauskiene et al., 2015). However, the responsiveness of spliceosome under hypoxia treatment remains to be elucidated *in planta*. In this study, a variety of splicing components have been identified to show differential expression under hypoxia treatment. Among these, six isoforms from two SR proteins (Os03g0344100, *SR32* and Os02g0610600, *RSZ23*) were induced by hypoxia treatment (Fig. 7E). Although multiple isoforms of SR proteins have been detected in different rice tissues (Peng et al., 2013), no evidence links them to hypoxia stress responsiveness before. Here, we hypothesize that SF changes under hypoxia is crucial for downstream AS regulation under hypoxia. However, less information can be found by annotation and datamining of these SFs that we have identified in this study. Further functional characterization is required to confirm their roles in response to hypoxia. Besides post-transcriptional regulatory pathways, processes related to protein export, lysosome and proteasome were observed to play a role during hypoxic germination (Figs. 1 and 2). The enhancement of some splicing isoforms in protein export process (Fig. 2C) may effectively help plants to survive during hypoxia conditions. Furthermore, lysosome is a place where cell to recycle building materials or detoxification (Chen et al., 2015). Recent study shows that hypoxia may rapidly induce autophagy, which is a highly conserved mechanism in eukaryotes to target cellular components to lysosome for recycling purpose (Chen et al., 2015). Thus, the newly formed isoforms of lysosomal gene may be responsible for the survival under hypoxia stress. Similarly, protein degradation has been considered as a major responsive mechanism in response to hypoxia in both animals and plants (Huang et al., 1998; Gibbs et al., 2011; Licausi et al., 2011). Significant protein will be degraded as an alternative energy source and remodelling during hypoxia treatment. Isoforms formed in this process may efficiently degrade misfolded proteins for synthesis of proteins isoforms that can confer hypoxia tolerance. Intriguingly, transcriptional regulation focused on the control of cellular metabolic levels and growth factors, whereas alternative splicing aims to produce new protein isoforms that is mainly involved in degradation, post-transcriptional regulation and transport processes. These two complementary mechanisms may facilitate rice seeds to survive under hypoxia during germination.

### Thousands of novel proteins or peptides resulting from alternative translation participate in the hypoxia response during rice germination

In addition to AS-resulting protein diversity, proteins encoded from a second frame of the same transcript or from annotated non-coding regions contribute to genome coding ability as well (Jensen et al., 2013; Wade and Grainger, 2014). Specifically, a considerable number of unannotated proteins were detected using a customized library by six-frame translation (*i.e.*, 3 in the forward strand and 3 in the reverse complement strand). The coding ability of one transcript using a second frame has been widely studied in animals but is rarely reported in plants (de Klerk and t Hoen, 2015). One example from plant systems is an alpha-enolase gene (*LOS2*) in Arabidopsis that encodes an MBP-like protein by alternative translation. This MBP-like protein affects ABA responses and its protein level is regulated by E3 ligase SAP5 (Kang et al., 2013). Furthermore, the existence of uORFs in the 5’-untranslated regions of certain transcripts may lead to a feedback regulation of translation efficiency (Laing et al., 2015). From our results, a total of 2,660 putative proteins over 80 amino acids and 904 proteins/peptides ranging from 6 to 80 amino acids have been identified (Fig. 5D). A total of 960 of these proteins/peptides were specifically induced under hypoxia treatment, suggesting that they are new players involved in hypoxia responses. Furthermore, a total of 137 novel proteins were quantified by proteomic analysis, 128 of which were not present in the DEG and DEP lists (Fig. 5E), demonstrating that the usage of a customized library combined with quantitative proteomics is essential for this kind of novel protein/peptide identification.

### Proteogenomic approach evolves as a new generation method to analyse omics-based datasets

Large profiling methods have been applied in plant research to study various developmental processes or stress responses. However, individual approaches such as transcriptome or proteome analysis are restricted by their defects in experimental conditions and analytical pipelines. For example, pure transcriptome analysis is affected by the corresponding reference genome annotation. Pure proteomic methods are limited by currently available protein libraries, which were generated based on incomplete genome information (Zhu et al., 2017). Thus, proteogenomics, a method incorporating transcriptomic and proteomic datasets, represents a new generation of analytical approaches for deeper understanding of the functional importance of potential genome coding ability (Castellana et al., 2008; Kumar et al., 2016). First, this analytical approach is able to determine which AS isoforms will be translated into proteins and thus can differentiate between mRNA degradation regulation and translational control (Nicholson et al., 2010; Drechsel et al., 2013). Second, in combination with quantitative proteomics, proteogenomic analysis links the protein evidence to their transcript changes to give an accurate footprint for each transcript isoform during the analysis. Low correlation of the expression levels between proteins and transcripts will be improved when using this type of analytical pipeline (Fig. 4D, E). This in turn will reveal valuable targets that are truly regulated at transcript and protein levels in the same trend. At last, coupled with a self-constructed protein library, this method enhances the identification of novel proteins/peptides (Fig. 5) that are potential hidden regulatory components in plant development or stress responses. However, this approach can be further improved from its current version. For example, using strand-specific library construction in short-read RNA seq analysis can enhance the accuracy and reduce redundancy of subsequent protein library construction. Furthermore, using the 3^rd^ generation of sequencing methods, such as single molecule long-read sequencing, can aid in the precise identification of full-length transcripts for accurate AS identification (Zhu et al., 2017). In addition, the low overlap between DEG and DEP or DAS and DASP can be improved by increasing the throughput and coverage of proteomic analysis. The incorporation of SWATH (sequential window acquisition of all theoretical spectra-mass spectrometry)-based quantitative proteomics (Zhu et al., 2016; Zhu et al., 2016) and two or more enzyme digestion steps may achieve better results than those of the current study.

## CONCLUSION

In conclusion, this study expands our understanding of the genome coding ability of rice under hypoxic germination. Two post-transcriptional mechanisms, alternative splicing and alternative translation initiation, have major contributions to protein diversity during hypoxia (Figure 8). Alternative splicing may function in parallel with transcriptional control in response to hypoxia stress during rice germination. Specifically, low oxygen conditions extensively affect AS and ATI patterns in parallel with conventional transcriptional regulation during rice germination. The compositional change of spliceosomes may result in the preferred usage of non-canonical splicing sites under hypoxia treatment. In this case, the conservation of 5’-splicing sites was largely affected by the hypoxia treatment. In addition, hypoxia-affected DAS events were more likely undergo protein translation in comparison to AS events identified under normal conditions. The above results indicate the existence of a large underground network of hypoxia responses at the post-transcriptional level. This newly discovered underlying response mechanism is mediated by AS and ATI. The members of this network need to be further characterized. This case study using hypoxic germination as a model demonstrates how modern technology and bioinformatic analysis improves our understanding of the plant genome coding ability and its features during stress responses.

**Figure 8.**
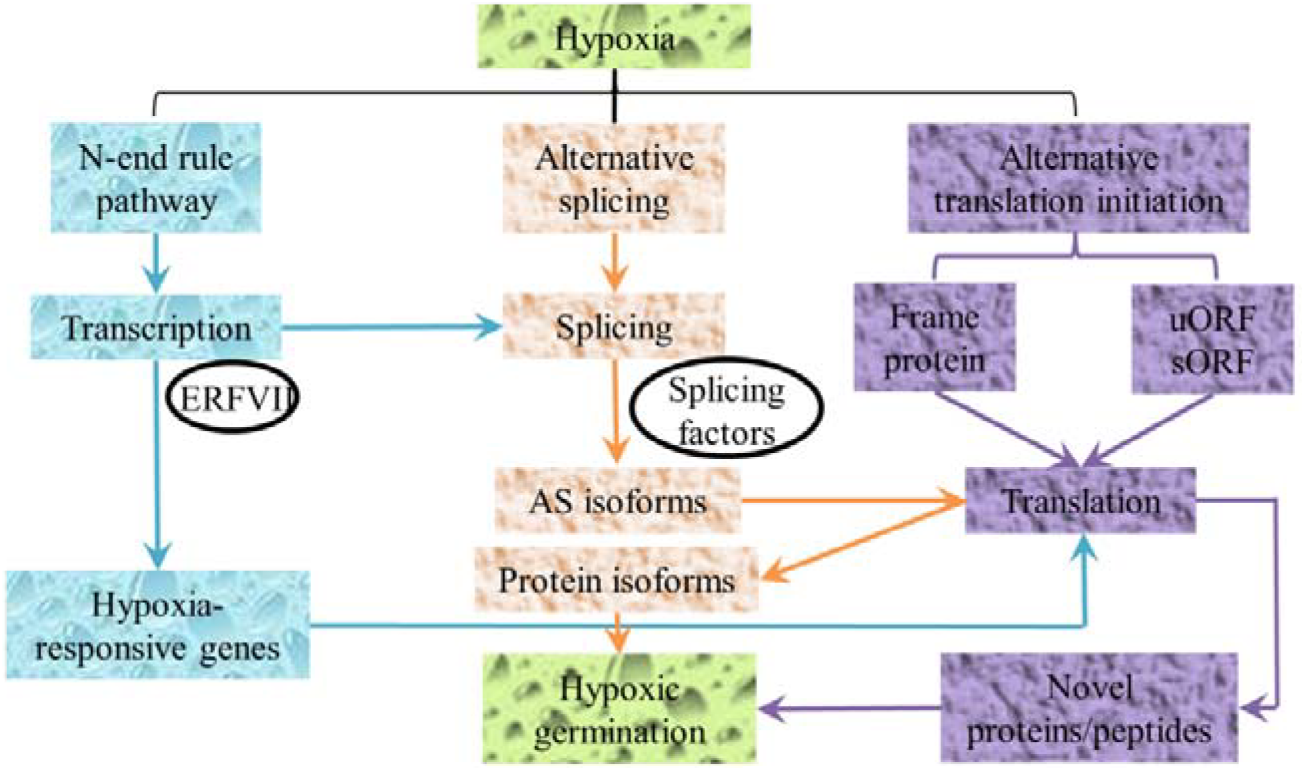
Model of alternative splicing and alternative translation initiation involved in the hypoxic germination pathway. Summary model of the rice genome using its coding ability to produce diverse functional proteins during hypoxic germination. The traditional transcriptional pathway (blue) has been well studied. The parallel pathway of alternative splicing (AS, orange) is able to generate AS isoforms, which in turn can be translated into protein isoforms in response to hypoxia treatment. In the third pathway of alternative translation initiation (ATI, violet), upstream open reading frames (uORFs) and small ORFs (sORFs) can further expand the protein diversity under hypoxia treatment.

## Supplemental Data

**Supplemental Fig. 1.** Analytical pipeline of AS identification, quantification and validation in this study.

**Supplemental Fig. 2.** Comparison of previous published datasets and qRT PCR validation.

**Supplemental Fig. 3.** GO enrichment analysis between DAS and DEG datasets from RNA sequencing.

**Supplemental Fig. 4.** qRT-PCR validation of selected genes from DAS events.

**Supplemental Fig. 5.** Comparison of splicing sites (ss) recognition between AS and DAS events.

**Supplemental Table 1.** Summary of the basic parameters in RNA sequencing dataset.

**Supplemental Table 2.** List of differentially expressed genes.

**Supplemental Table 3.** List of the differentially expressed AS events.

**Supplemental Table 4.** Summary of quantified proteins in proteomic analysis.

**Supplemental Table 5.** Primers used in this study.

## ACKNOWLEDGMENTS

This work was supported by the Natural Science Foundation of Shandong Province (BS2015NY002), Funds of Shandong “Double Top” Program, Natural Science Foundation of China (NSFC31101099), the China Postdoctoral Science Foundation (2017M622801), Science and Technology Program of Nantong (MS12016044), the National Natural Science Foundation of China (NSFC31101099, 31771701, 31701341), Innovative Training Program of Nantong University College Students 2017 (201710304049Z), National Key Basic Research Program of China (2012CB114300), the Natural Science Foundation of Guangdong Province (2014A030313794), Shenzhen Overseas Talents Innovation and Entrepreneurship Funding Scheme (The Peacock Scheme, KQTD201101) and Hong Kong Research Grant Council (AoE/M-05/12, AoE/M-403/16, CUHK 14122415, 14160516, 14177617).

## AUTHOR CONTRIBUTIONS

M.X.C., F.Y.Z., J.H.Z., Y.G.L. designed experiments. M.X.C., F.Y.Z., F.Z.W., N.H.Y., T.F., Y.Y.C., T.Y.L. S.S.Z performed experiments. M.X.C., F.Y.Z., B.G., K.L.M., G.Y.F., Z.Z.S., L.J.X., Q.J.H., H.J.W. analysed data. F.Y.Z., M.X.C., N.H.Y. wrote the manuscript. S.X., J.H.Z., Y.G.L. critically commented and revised it.

---

**Figure S1 Analytical pipeline of AS identification, quantification and validation in this study.**

**Figure S2 Comparison of previous published datasets and qRT PCR validation.**

(A) Heatmap comparison of previous published microarray datasets (Pub) to our RNA seq analysis (Our). (B) qRT-PCR validations of the selected and marker genes during hypoxic germination from three biological replicates. *OsACTIN1* was used as an internal reference gene. ‘*’ and ‘**’ denote that the relative mRNA level is significantly higher in hypoxia-treated samples (grey bars) in comparison to air control (black bars) in complete darkness, *P*<0.05 and *P*<0.01, respectively. **Locus IDs (Bolded)** represent genes have similar expression pattern in qRT-PCR analysis in comparison to previous transcriptome or microarray analysis.

**Figure S3 GO enrichment analysis between DAS and DEG datasets from RNA sequencing.**

**Figure S4 qRT-PCR validation of selected genes from DAS events.** Primers used in the experiment are listed in Supplemental Table 5. *OsACTIN1*was used as an internal reference gene. ‘*’ and ‘**’ denote that the relative mRNA level is significantly higher or lower in hypoxia-treated samples in comparison to air control, *P*<0.05 and *P*<0.01, respectively. AS events in bold form represent the consistency between RNA seq and qRT-PCR data.

**Figure S5 Comparison of splicing sites (ss) recognition between AS and DAS events.** ALE, alternative last exon; AFE, alternative first exon; SKIP, exon skipping; IR, intron retention; AE5’, alternative donor; AE3’, alternative acceptor.

